# “Sifarchaeota” a novel Asgard phylum capable of polysaccharide degradation and anaerobic methylotrophy

**DOI:** 10.1101/2020.10.14.339440

**Authors:** Ibrahim F. Farag, Rui Zhao, Jennifer F. Biddle

## Abstract

The Asgard superphylum is a deeply branching monophyletic group of Archaea, recently described as some of the closest relatives of the eukaryotic ancestor. The wide application of genomic analyses from metagenome sequencing has established six distinct phyla, whose genomes encode for diverse metabolic capacities and play important biogeochemical and ecological roles in marine sediments. Here, we describe two metagenome-assembled genomes (MAGs) recovered from deep marine sediments off Costa Rica margin, defining a novel lineage phylogenetically married to Thorarchaeota, as such we propose the name “Sifarchaeota” for this phylum. The two “Sifarchaeota” MAGs encode for an anaerobic methylotrophy pathway enabling the utilization of C1-C3 compounds (methanol and methylamines) to synthesize acetyl CoA. Also, the MAGs showed a remarkable saccharolytic capabilities compared to other Asgard lineages and encoded for diverse classes of carbohydrate active enzymes (CAZymes) targeting different mono-, di- and oligosaccharides. Comparative genomic analysis based on the full metabolic profiles of Asgard lineages revealed the close relation between “Sifarchaeota” and Odinarchaeota MAGs, which suggested a similar metabolic potentials and ecological roles. Furthermore, we identified multiple potential horizontal gene transfer (HGT) events from different bacterial donors within “Sifarchaetoa” MAGs, which hypothetically expanded “Sifarchaeota” capacities for substrate utilization, energy production and niche adaptation.

**Importance:** Deep marine sediments are the home of multiple poorly described archaeal lineages, many of which have ecological and evolutionary importance. We recovered metagenome-assembled genomes (MAGs) belonging to a novel Asgard phylum from the deep sediment of the Costa Rica margin. We proposed the name “Sifarchaeota” to describe the members of this phylum. Representative genomes of the “Sifarchaeota” showed remarkable saccharolytic capacities extending the known metabolic features encoded by the Asgard lineages. We attribute its ability to survive under the deep sediment conditions to its putative capacities to utilize different (C1-C3) compounds commonly encountered in deep sediment environments via anaerobic methylotrophy pathway. Also, we showed the importance of horizontal gene transfer in enhancing the “Sifarchaeota” collective adaptation strategies.

## Introduction

Deep marine sediments are the home of multiple poorly described archaeal lineages, most of which are yet uncultured (1)(2). Recently, the discovery of Asgard archaea in benthic environments has generated great interest in novel lineages from marine sediments. Additionally, greater attention directed towards studying Asgard archaea is also in part to understanding eukaryogenesis, as this superphylum harbors the most closely related archaeal group to eukaryotes and their genomes encode for multiple homologs of eukaryotic proteins (3)(4). So far, there are 6 established Asgard phyla: Lokiarchaeota, Thorarchaeota, Odinarchaeota, Heimdallarchaeota, Helarchaeota and Gerdarchaeota (4)(5)(6). The number of novel lineages yet to be found is at this point unknown. This raises the need for more genome resolved metagenome surveys to recover genomes of these archaeal lineages, decipher their metabolic capacities and place them in the context of microbial ecology. Previous studies targeting the deep sediment from the Costa Rica margin subseafloor have shown the presence of diverse archaeal communities (7)(8). Among these archaeal lineages, members of Asgard superphyla were highly abundant at multiple depths, making up 17% of the archaeal communities present (7)(8). In this study on the Costa Rica margin subseafloor, we employ genome resolved metagenomics to describe the metabolic potential of the genomes belonging to a new Asgard phylum. Phylogenomic analysis placed the sequences of the new Asgard genomes as a sister clade to the sequences of Thorarchaeota and we propose the name “Sifarchaeota” to describe this new Asgard phylum. Comparative analysis shows distinct differences between Sifarchaeota and previously reported Asgard archaea in terms of substrate utilization, energy production and niche adaptation strategies. Finally, we detect multiple potential horizontal gene transfer (HGT) events from different bacterial donors expanding the substrate utilization, energy production and secondary metabolite production capacities of the Sifarchaeota.

## Methods

### Sample collection

Samples were collected during International Ocean Drilling Program (IODP) Expedition 334, Site U1379B on the Costa Rican Margin. Details on sample location and sampling methodology have been previously described (7)(9). Microbiology samples (whole-round cores) were collected on board and frozen immediately at −80°C. They were shipped to the Gulf Core Repository (College Station, Texas) on dry ice and stored at −80°C until shipping to the Biddle lab (Lewes, Delaware) on dry ice and further storage at −80°C. Metagenome sequencing data were generated from four silty clay sediment horizons (2H-1, 2H-2, 2H-5A, and 2H-5B) in the depth interval of 2-9 mbsf, within the sulfate reduction zone (7).

### DNA extraction, library construction and sequencing

DNA for metagenomic sequencing was extracted from ~7 g sediment (~0.7 g sediment in 10 individual lysis tubes) using PowerSoil DNA Isolation Kit (Qiagen) following the manufacturer’s instructions, except for the following minor modification: the lysing tubes were incubated in water bath of 60°C for 15 min prior to beading beating on a MP machine at speed 6 for 45 seconds. The DNA extracts were iteratively eluted from the 10 spin columns into a final of 100 μL of double distilled H2O for further analysis. Metagenomic libraries were prepared and sequenced (150 bp paired-end) on an Illumina NextSeq 500 sequencer at the Genome Sequencing & Genotype Center at the University of Delaware.

### Assembly and genome binning

The raw sequencing data were processed with Trimmomatic v.0.36 (10) to remove Illumina adapters and low quality reads (“SLIDINGWINDOW:10:25”). The quality-controlled reads from the eight samples were de novo co-assembled into contigs using Megahit v.1.1.2 (11) with the k-mer length varying from 27 to 117. Contigs longer than 1000 bp were automatically binned using MaxBin2 (12) and Metabat2 (13), and the best quality ones were selected using DAS_Tool (14) with the default parameters. The resulting MAGs were quality assessed using CheckM (15) and taxonomically classified using GTDBTk v1.3.0 (16) using the default parameters. Genome bins of >50% completeness were manually refined using the gbtools (17) based on the GC content, taxonomic assignments, and differential coverages in different samples. Coverages of contigs in each sample were determined by mapping trimmed reads onto the contigs using BBMap v.37.61 (18). Taxonomy of contigs were assigned according to the taxonomy of the single-copy marker genes in contigs identified using a script modified from blobology (19) and classified by BLASTn. SSU rRNA sequences in contigs were identified using Barrnap (Seeman 2015, Github), and classified using VSEARCH with the SILVA 132 release (20) as the reference.

To improve the quality of the two novel Asgard archaea MAGs, we recruited quality-controlled reads using BBMap from 2H-2, because the highest genome coverages of these two MAGs were detected in this particular sample. The recruited reads were then re-assembled using SPAdes v.3.12.0 (21) using default parameters. After removal of contigs shorter than 1 kb, the resulting scaffolds were visualized and re-binned manually using gbtools (17) as described above. The quality of the resulting Asgard archaea genomes were checked using CheckM v.1.0.7 (15) with the “lineage_wf” option.

### Concatenated ribosomal protein phylogeny

To determine the phylogenetic affiliations of the two Sifarchaeota MAGs in the Archaea domain, we performed a thorough phylogenomic analysis based on the concatenation of 16 ribosomal proteins (L2, L3, L4, L5, L6, L14, L15, L16, L18, L22, L24, S3, S8, S10, S17, and S19). Reference genomes were selected from all the major archaeal phyla (3-5 for each) included in the GTDB database (22), except for the Asgard superphylum for which all available genomes were included. Ribosomal protein sequences were detected in Anvi’o (23) using the respective HMM profiles, aligned using MUSCLE v3.8.31(24), and concatenated. The maximum likelihood phylogenetic tree was reconstructed using IQ-Tree (v1.6.6) (25) (located on the CIPRES web server) (26) with VT + F + R10 as the best-fit substitution model selected by ModelFinder (27), and single branch location was tested using 1000 ultrafast bootstraps and approximate Bayesian computation (28). Branches with bootstrap support >80% were marked by black circles. In addition to phylogenomic analysis, we also calculated the average nucleotide identity (ANI) using FastANI (29) and average amino acid identity (AAI) using CompareM (https://github.com/dparks1134/CompareM) with default settings between these novel Asgard MAGs and other public available ones (i.e., those included in the GTDB database), to further explore the novelty of these MAGs.

### Metabolic reconstruction

Amino acid sequences encoded by the Sifarchaeota MAGs were predicted using Prodigal v2.6.3 (30) applying the default parameters and using translation table 11. The resulting amino acid sequences were screened using HMMsearch tool (31) against custom HMM databases (32) representing the key genes for specific metabolic pathways to understand the potential metabolic capacities and their ecological roles of Sifarchaeota. The presence/absence profiles of the metabolic pathways and their completion levels were further assessed through querying the predicted amino acids against KEGG database using BlastKoala tool (33). Carbohydrate-active enzymes encoded by the Sifarchaeota MAGs were analyzed using dbCAN-fam-HMMs (v6) database (34). Proteases, peptidases, and peptidase inhibitors encoded by the MAGS were detected via USEARCH-ublast tool (35) against the MEROPS database v12.1(36). Finally, the predicted amino acid sequences were queried against the TCDB database (37) using USEARCH-ublast tool (35) to identify the potential transporters.

### Genome centric comparative analysis for the Asgard superphylum

We compared the total metabolic profiles of Sifarchaetoa MAGs with representatives from other Asgard lineages including (Odinarchaetoa, Thorarchaeota, Lokiarchaeota and Heimdallarcaheota) to identify the key metabolic differences between each of the phyla within the Asgard superphylum. We queried the representative MAGs of each of the Asgard lineages against KOfam database (KEGG release 94.1) via KofamKOALA webtool (38) and using evalue 10^-3^. Then, the Asgard MAGs were clustered based on the presence/absence profiles of the identified KOfams, shared between at least 3 genomes, using clustergrammer web-based tool (39) and applying the Euclidean distance and average linkage type.

### Horizontal gene transfer (HGT) analysis

The HGT events were detected through querying Sifarchaeota predicted amino acids against KEGG database (KEGG release 94.1) via GhostKoala web interface (33). Proteins with KO annotation, non-Asgard hits, and a bit score >100 were considered as potential HGT candidate proteins. Then, the candidate set of proteins were queried against (nr) and UniProtKB/swissprot databases and all proteins showing Asgard hits as one of the top hits were removed from any further analysis. Finally, each candidate protein is aligned to reference set of proteins collected via Annotree (40) using the corresponding KO entry and the HGT events were confirmed through creating approximately-maximum-likelihood phylogenetic tree for each candidate protein using FastTree v2.1 (41). An outline for the HGT detection pipeline is illustrated in Supplementary Figure 1.

## Results

### MAG construction and phylogenomic analysis

We reconstructed 3 MAGs belonging to a potentially novel Asgard phylum with moderate completion levels (67-80%) and very low contamination levels (0.93-1.9%) (Table 1). The taxonomic affiliations of these potentially novel Asgard MAGs were tested using 16 ribosomal proteins phylogenomic tree and it showed that both MAGs clustered together and formed a sister lineage to MAGs belonging to the phylum Thorarchaeota (Figure 1). The phylogenetic analysis using a set of 122 archaea specific marker proteins implemented in GTDB-tk confirmed that these MAGs are belonging to a potentially novel Asgard archaeal lineage. To confirm the unique positions of these candidate novel Asgard lineage, we calculated the average nucleotide identities (AAI) and average amino acid identities (ANI) between the 3 novel Asgard MAGs and other MAGs representing the other Asgard lineages including (Lokiarchaeota, Thorarchaeota, Heimdallarchaeota and Odinarchaeota). On average, the 3 novel Asgard MAGs showed low AAI values when compared to the other Asgard MAGs (<50%) (Supplementary Tables 1 and 2).

**Figure 1.**
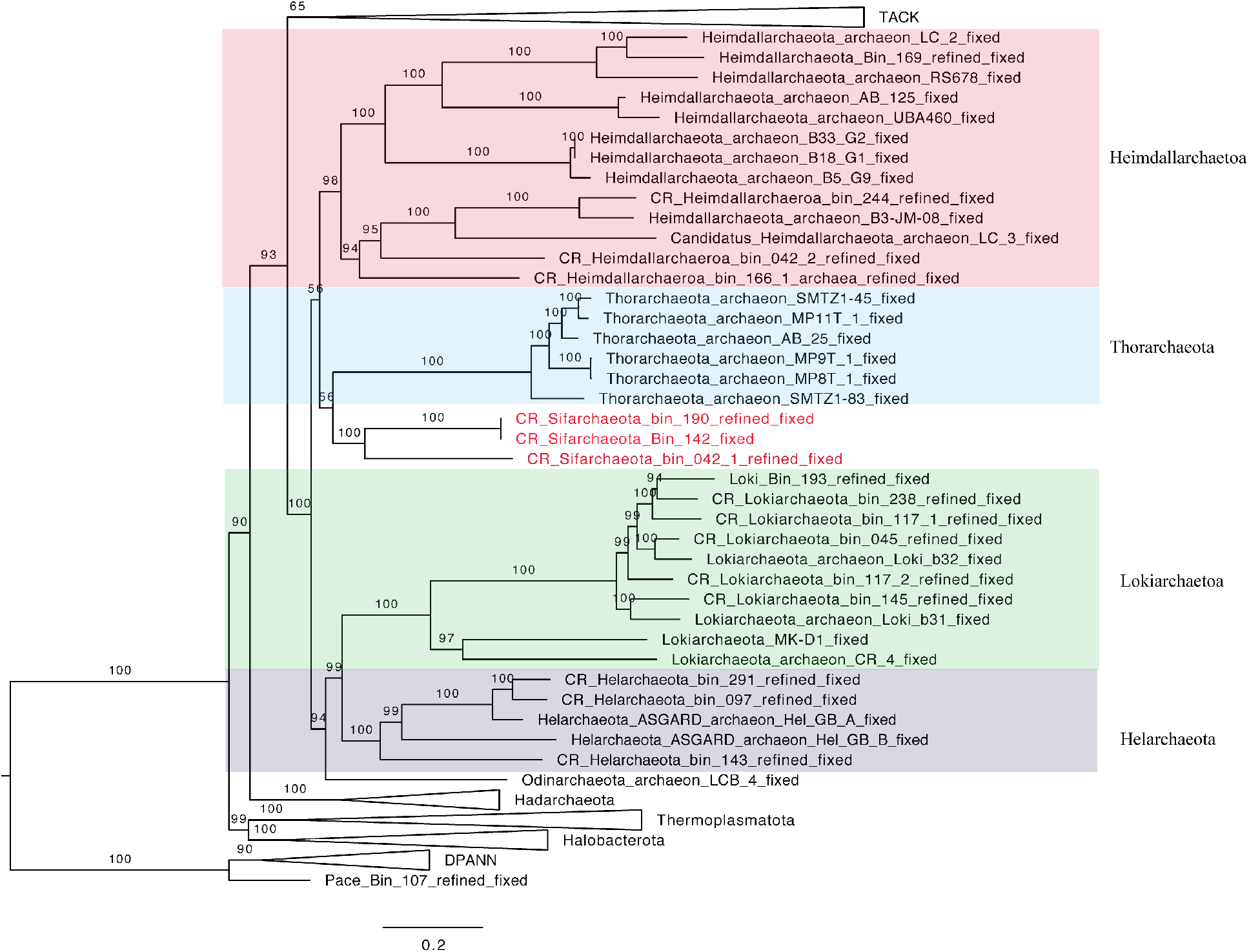
Maximum-likelihood phylogenetic tree of archaea genomes based on concatenated 16 ribosomal proteins. This tree was inferred using IQ-TREE v1.6.10 with the LG+R7 model and 1000 ultrafast bootstraps. The Sifarchaeota MAGs recovered in this study is highlighted in red. Lineages of the Asgard superphylum are expanded, while the other linages were collapsed, if possible. The scale bar shows estimated sequence substitutions per residue.

**Figure 2.**
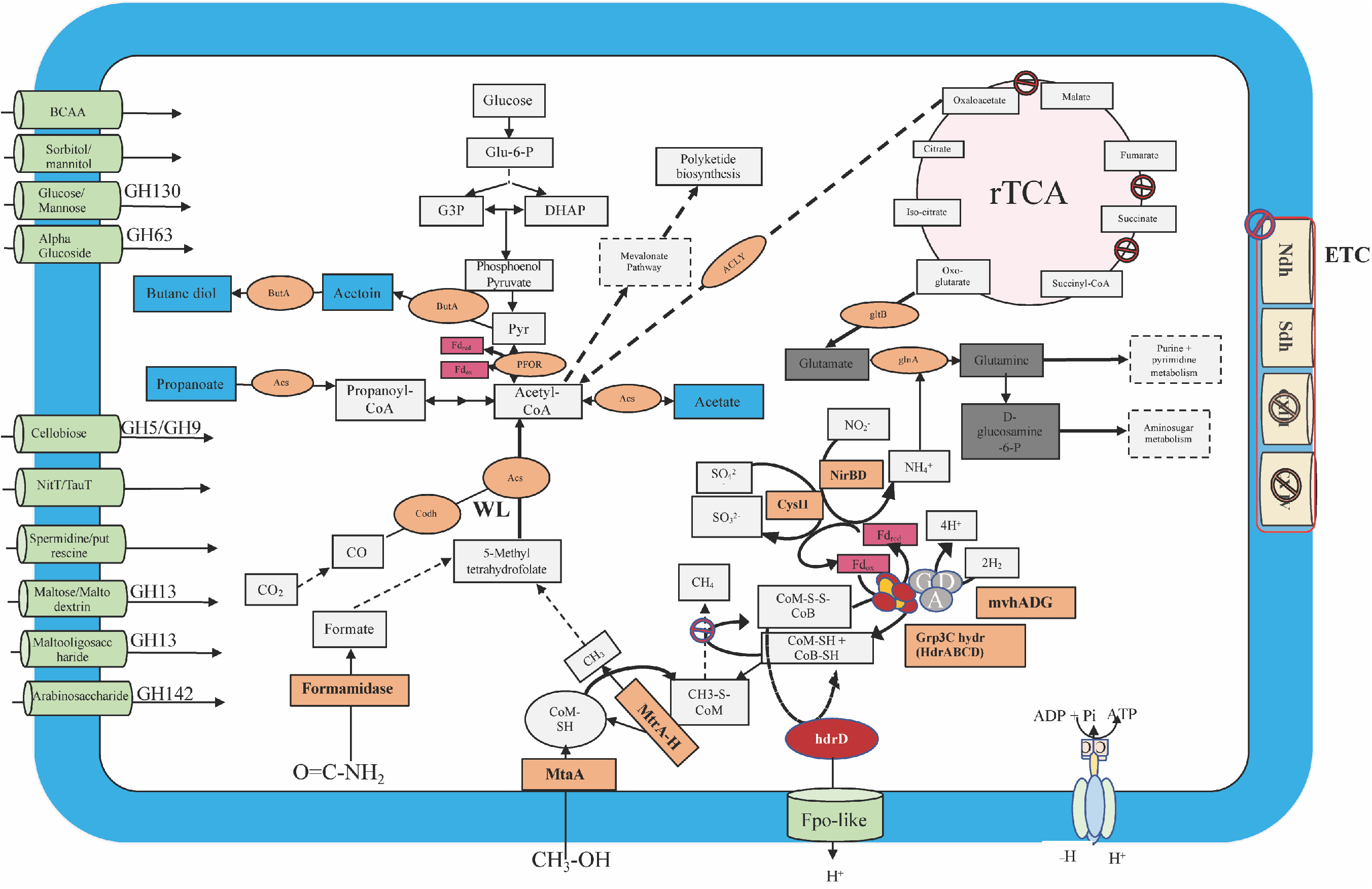
Metabolic reconstruction of the key metabolic pathways encoded by the Sifarchaeota MAGs. Central metabolic pathways are shown in gray boxes, carbon fixation pathways (WL and rTCA cycles) are shown in pink, electron transport chain (ETC) proteins are shown in yellow, fermentation products are shown in blue boxes, enzymes and enzyme complexes are shown in orange circles, energy carriers are shown in red, and metabolite and amino acid transporters are shown in light green.

**Figure 3.**
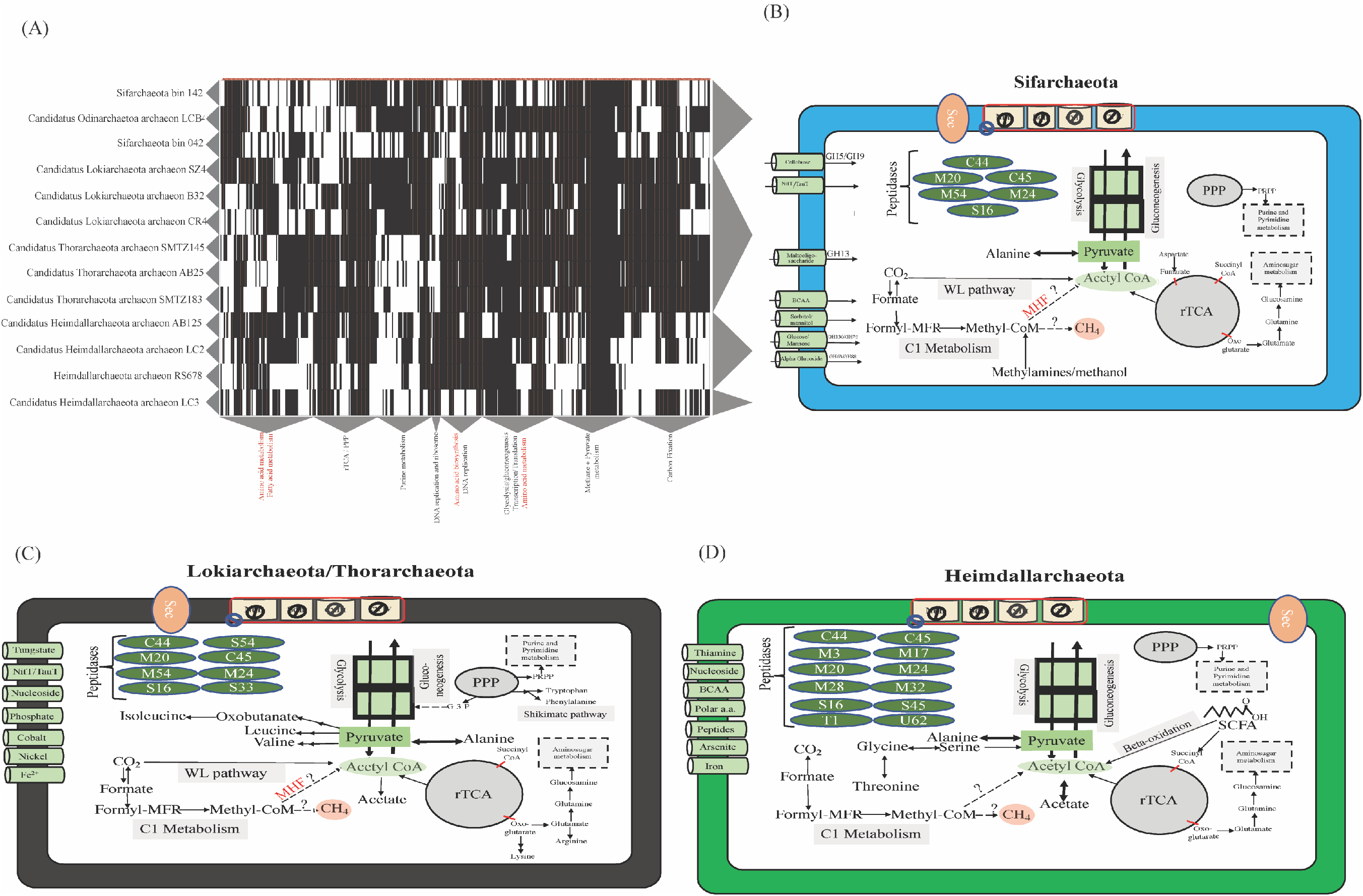
Comparative analysis between MAGs representing different Asgard members. (A) Heatmap clustering the different Asgard MAGs (X-axis) based on their total metabolic profiles as predicted by Kofam database (Y-axis). The clustering was performed using Euclidean distance and complete linkage methods. (B) Metabolic model illustrates the key metabolic features identified in Sifarchaeota cluster. (C) Metabolic model illustrates the key metabolic features identified in Lokiarchaeota and Thorarchaeota cluster. (D) Metabolic model illustrates the key metabolic features identified in Heimdallarchaeota clusters.

**Table 1.** Details of the Sifarchaeota MAGs analyzed in this study.

| Bin ID | Completion | Contamination | Genome Size (Mbps) | Estimated Genome Size (Mbps) | Strain heterogeneity | # contigs |
| --- | --- | --- | --- | --- | --- | --- |
| Bin.042_1 | 67.76 | 0.93 | 1.8 | 2.39 | 0 | 301 |
| Bin_190 | 74.4 | 1.9 | 1.9 | 2.38 | 0 | 786 |
| Bin_142 | 81.4 | 1.9 | 2.01 | 2.39 | 0 | 300 |

Two of the new Asgard MAGs, bins 190 and 142, showed a very high similarity with AAI 98.81% and ANI 99.52%, as such, we focused all our subsequent analysis on two MAGs, bins 042 and 142, since they were the most complete and unique MAGs in this study.

### Description of the taxa

Although the two Sifarchaeota MAGs, bin042 and bin142, shared many metabolic similarities, we could not describe them using one type strain due to the significant phylogenetic distances between the two MAGs (Figure 1) and the genomic differences as described using ANI and AAI values (Supplementary Tables 1 and 2). Therefore, we used two type strains to describe the two putative Sifarchaeota lineages.

*Candidatus* Sifarchaeotum costaricensis (***costaricensis* of or from Costa Rica**). Type material is the genome designated as bin042 representing ‘*Candidatus* Sifarchaeotum costaricensis’.

*Candidatus* Sifarchaeotum subterraneus (***subterraneus* latin name of subsurface**). Type material is the genome designated as bin142 representing *‘Candidatus* Sifarchaeotum subterraneus’.

Based on these genera, we further propose the name of a new Asgard phylum, the phylum ‘*Candidatus* Sifarchaeota phylum nov.’

### Metabolic reconstruction of Sifarchaeota MAGs showed remarkable saccharolytic capacities and potential anaerobic methylotrophy

Previous reports showed that Asgard genomes encode for wide range of protein and fatty acids degradation capacities (42). So far, most of the known Asgard genomes showed limited saccharolytic capacities emphasized by their low genomic densities of CAZymes and sugar transporters. However, Sifarchaeota showed high abundance and diversity of <u>C</u>arbohydrate <u>A</u>ctive <u>E</u>nzymes (CAZymes) encoded by their MAGs specifically targeting sugars varying in complexities from low (C1-C3) to moderate (C4-C6), including mono-, di- and oligo-saccharides (Supplementary Figure 1, Supplementary Tables 3 and 5). Interestingly, CAZyme analysis revealed the presence of different glycoside hydrolases (GH) families including cellulases and endoglucanases (GH5 and GH9), cyclomaltodextrinase (GH13), α-glucosidase (GH63), β-1,4-mannooligosaccharide phosphorylase (GH130) and β-L-arabinofuranosidase (GH142) targeting cellobiose, maltose and maltooligosaccharides, alpha-glucosaccharides, glucose/mannose and arabinosaccharides, respectively. We also identified dedicated sugar transporters mediating the transfer of these sugars inside the Sifarchaeota cells, where the degradation and fermentation processes take place (Supplementary Table 6). Metabolic reconstruction of Sifarchaeota MAGs predict a general fermentative anaerobic life style with multiple anaerobic respiration capabilities emphasized by the presence of incomplete reverse TCA cycle, presence of various fermentation pathways capable of producing different fermentative products including acetate, acetoin and butanediol under strictly anaerobic conditions as well as the potential capabilities to anaerobically respire sulfate to sulfite and nitrite to ammonia. Notably, the degradation capacities for proteins, peptides and amino acids as well as amino acid metabolism are limited in Sifarchaeota compared to the other benthic archaea. Hence, Sifarchaeota may make up the short supply of fixed nitrogen via reducing nitrite to ammonia and encoding for amidases to extract fixed nitrogen from nitrogen containing amides like formamide.

Sifarchaeota MAGs showed the capacity to utilize various C1 compounds including formate and methanol. For example, methanol is metabolized using an anaerobic methylotrophy pathway previously described (8), suggesting the potential widespread distribution of this pathway among benthic marine archaea to metabolize methylated compounds. This potential anaerobic methylotrophic capacity was inferred via detecting incomplete methylotrophic methanogenesis pathway, where the key genes encoding for the methyl coenzyme reductase complex were completely absent. Besides, genes for incomplete Wood Ljundahl (WL) pathway were identified, where only the genes of the carbonyl branch were present and the genes of the methyl branch were completely absent. Detecting these set of genes in Sifarchaeota MAGs suggests the presence of anaerobic methylotrophic capability, enabling Sifarchaeota to recycle methyl groups within methanol and other methylated compounds. Then, these methyl groups are transferred to tetrahydrofolate complex replacing the function of the methyl branch of WL pathway and ultimately producing acetyl CoA.

### Comparative genomic analyses between different Asgard lineages shows diverse metabolic features and life style patterns

To understand the key metabolic differences between the Asgard lineages, we conducted a comprehensive genome-centric analyses. As described in the methods section, we parsed 13 different MAGs (2 MAGs obtained from this study and 11 publicly available MAGs) belonging to 5 different Asgard phyla (Lokiarchaeota, Thorarchaeota, Heimdallarchaeota, Odinarchaeota and Sifarchaeota) against KOfam database using an HMM search tool (Supplementary Table 7. The analysis output focused on two aspects (1) exploring the range of life style diversity within the Asgard superphylum and (2) comparing the metabolic capabilities of different Asgard lineages.

Overall, the comparative analyses grouped the Asgard superphylum into 4 distinct clusters based on their whole metabolic profiles. Notably, all the MAGs within the Asgard superphylum suggested a similar fermentative anaerobic life styles, where the core genes for reverse TCA cycle, various fermentation pathways were present as well as the absence of oxidative phosphorylation and oxygen tolerance related genes. Here we describe the clusters and the features that drive their clustering.

The **first cluster** grouped the 2 Sifarchaeota MAGs (bin42 and 142), obtained from this study, with Candidatus Odinarchaetoa archaeon LCB4. Sifarchaeota MAGs shared multiple metabolic similarities with the Odinarchaeota MAGs including limited amino acid and fatty acids metabolic potentials, evident saccharolytic activities, emphasized by high density of CAZyme encoding genes targeting different mono-, di- and oligosaccharide. Also, we could only identify genes encoding for the oxidative branch of the pentose phosphate pathway, producing Phosphoribosyl pyrophosphate (PRPP), which eventually channeled to the purine and pyrimidine metabolic pathways to be used for nucleotide and nucleic acid biosynthesis. Also, MAGs within this cluster encoded for large number of genes involved in C1 metabolism including formate, methylamines and methanol. Genes encoding for formate dehydrogenase, methanol and methylamine specific corrinoid protein:coenzyme M methyltransferases were present, which suggest the potential capability of both lineages to utilize formate, methylamines and methanol as carbon sources, respectively.

The **second cluster** grouped Lokiarchaeota and Thorarchaeota MAGs together. Unlike the previous cluster, MAGs belonging to that cluster are characterized by their proteins and peptide degrading capabilities emphasized by the presence of high number of proteases and peptidases encoding genes ranging in density from (126-193 proteins/Mbs) and belonging to different families of serine and metalloproteases. Unlike other Asgard groups, Thorarchaeota and Lokiarchaeota showed the capacities to synthesize different amino acids including nonpolar amino acids (e.g. isoleucine, leucine, valine and alanine), aromatic amino acids (e.g. tryptophan and phenylalanine) via the shikimate pathway and charged amino acids (e.g. glutamate and lysine). Similar to other Asgard archaea, MAGs belonging to Thor- and Loki-archaeota encoded for genes mediating the metabolism of C1 compounds (e.g. formate), however, the absence of methyltransferases encoding genes excluded the use methylated compounds as one of the potential substrates.

Finally, **the third and fourth clusters** included different Heimdallarchaeota MAGs. The separation between Heimdallarchaeota based on their metabolic profiles suggests the presence of fundamental metabolic differences between the members of Heimdallarchaeota phylum and supports the previous findings that described Heimdallarcaheota as a polyphyletic group (3)(6). This raises the need for wider sampling efforts targeting Heimdallarchaeota genomes to fully resolve their phylogenetic position and evolutionary history. However, in this study we grouped all the Heimdallarchaeota MAGs and treated them as one phylum and designed a model that highlights the difference between them and the other Asgard lineages. Notably, all Heimdallarcahetoa MAGs showed the capacity to utilize proteins and short chain fatty acids as carbon sources, while polysaccharide degradation was less supported. Similar to Loki- and Throarchaeota MAGs, Heimdallarcahetoa MAGs encoded for high number of peptidases with coding densities (110-210 proteins/Mbs) and belonging to diverse families of proteases and peptidases including serine, metallo, cysteine and threonine peptidases (Supplementary Table 3).

Also, Heimdallarcahetoa MAGs showed the capacity to metabolize and synthesize nonpolar amino acids (e.g. alanine, glycine, and threonine). Among all the Asgard MAGs included in this analysis, only Heimdallarchaeota MAGs encoded for enzymes mediating beta-oxidation pathway including acyl CoA dehydrogenase, enoyl CoA hydratase and hydroxyacyl CoA dehydrogenase, suggesting their potential to use short-chain fatty acids (SCFA) as carbon and energy sources. Similar to all other Asgards, Heimdallarchaeota encoded for incomplete WL pathway, which could have a role in metabolizing C1 compounds like formate and formaldehyde.

Due to the limited access to the surrounding microbial community composition and environmental conditions as well as the underrepresentation of the some of the lineages (only 1 MAG from Odinarchaeota), we could not assess the exact reasons for these diverse substrate preferences between the Asgard phyla (e.g. polysaccharides in Sifarchaeota, proteins in Thor- and Loki-archaeota and short-chain fatty acids in Heimdallarchaeota). Also, we are unable to conclude at this point if these findings could be generalized for all the members within each phylum or it is just limited to the lineages/MAGs included in the study.

### Role of horizontal gene transfers (HGT) in enhancing Sifarchaeota metabolic capacities and niche adaptations

We investigated the role of HGT in expanding the metabolic capacities, substrate utilization and niche adaptation of the Sifarchaeota phylum. We traced the origin of each HGT event and identified the extent of the spread of each gene within the Asgard superphylum (Supplementary Figure 2). We successfully identified a total of 65 HGT events in Sifarchaeota, 12 (0.65% of the total proteins) and 53 (1.34% of the total proteins) events in bin042 and bin142, respectively. Most likely, the majority of the identified HGT events were lineage specific (58, 89.2%) and only few similar events were detected in other Asgard lineages (7, 11.8%). Also, we identified the potential donors for most of the horizontally transferred genes, ~90% of which were of bacterial origins. The major bacterial contributors for the horizontally transferred genes were Firmicutes (~30%), Chloroflexi (~15%), Proteobacteria (~13%), Cyanobacteria (5%) and other bacterial lineages (~30%). Only a small fraction (~10%) of the horizontally transferred genes were of archaeal origin outside of the Asgard superphylum (Figure 5 and Supplementary Table 8).

We classified HGT events based on the how widespread the transferred genes were among the different Asgard phylum, as well as other archaeal lineages. Accordingly, the HGT events were classified into lineage-specific, phylum-specific and domain-wide events. In the lineage specific event, the closest relatives of the transferred genes were bacteria and the genes were exclusively found in Sifarchaetoa MAGs and no orthologs were found in other Asgard or any other archaeal phyla (e.g. butanediol dehydrogenase). In the phylum-specific event, the closest relatives of the transferred genes were bacteria and the genes were found in multiple Asgard phyla (e.g. enoyl CoA hydratase). In the domain wide event, the closest relatives of the transferred genes were bacteria and the genes were found in different archaeal phyla (e.g. arsenate reductase *arsC* thioredoxin) (Figure 5). In general, the functional annotation of the transferred genes showed that these genes are involved in augmenting the Sifarchaeota metabolic repertoire. The majority of the transferred genes fall within two metabolic modules: butanoate metabolism and biosynthesis of secondary metabolites. In the butanoate metabolism, the majority of transferred genes were encoding for butane diol dehydrogenases and glutaconate CoA-transferase subunits mediating the key steps in pyruvate and acetate formation from butane diol and hydroxy glutaryl CoA, respectively.

**Figure 4.**
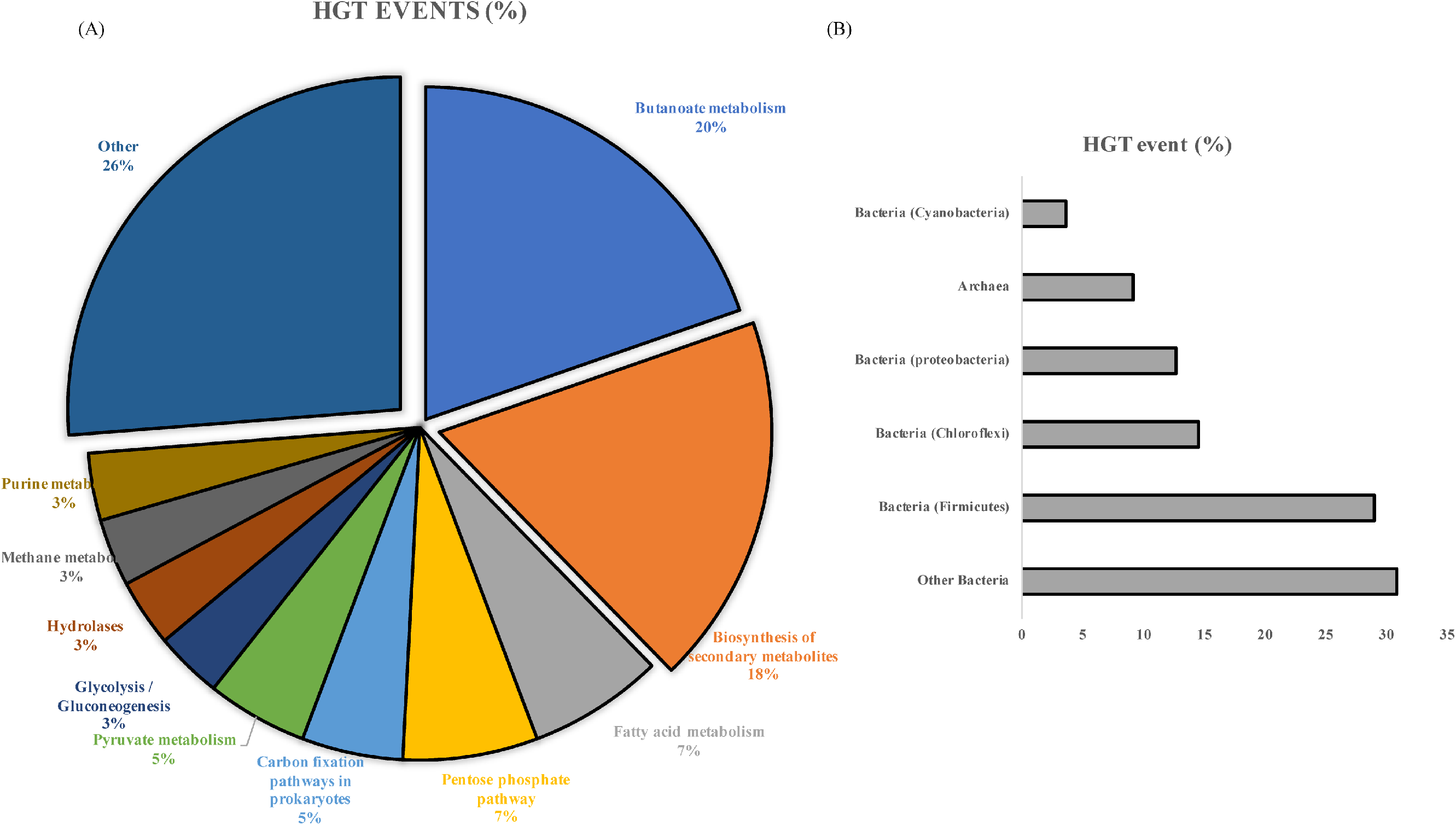
Summary of HGT events detected in the Sifarchaeota MAGs. (A) Different functional modules of horizontally transferred genes. (B) Major donors of the horizontally transferred genes.

**Figure 5.**
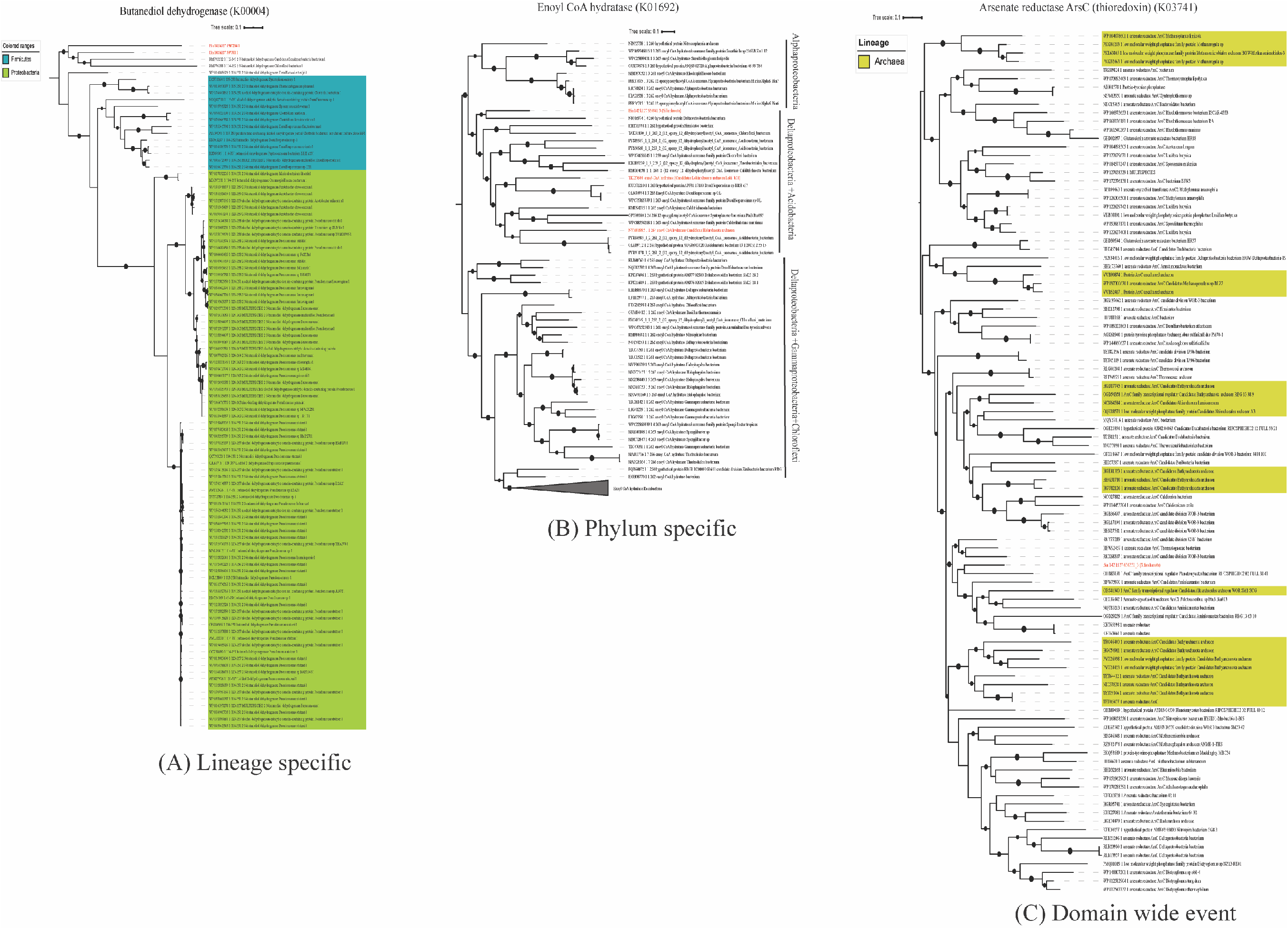
Examples for horizontally transferred genes in Sifarchaeota MAGs. (A) Lineage specific HGT event. Maximum likelihood phylogenetic tree of butanediol dehydrogenase gene sequences. Shaded areas correspond to the potential source organims. The tree was constructed on the basis of butanediol dehydrogenase gene sequences using FastTree. Reference sequences were obtained using AnnoTree (K00004). (B) Phylum specific HGT event. Maximum likelihood phylogenetic tree of enoyl CoA hydratase gene sequences. The tree was constructed on the basis of enoyl CoA hydratase gene sequences using FastTree. Reference sequences were obtained using AnnoTree (K01692). (C) Domain wide HGT event. Maximum likelihood phylogenetic tree of arsenate reductase ArsC thioredoxin gene sequences. The tree was constructed on the basis of arsenate reductase ArsC thioredoxin gene sequences using FastTree. Reference sequences were obtained using AnnoTree (K03741).

While the majority of the genes involved in the biosynthesis of secondary metabolites were part of the porphyrin and chlorophyll metabolism and terpenoid biosynthesis (e.g. anaerobic magnesium-protoporphyrin IX monomethyl ester cyclase and 1,4-dihydroxy-2-naphthoate polyprenyltransferase). Interestingly, multiple genes involved in niche adaptation may be acquired from the surrounding bacterial communities. These genes including formamidase and MtaA/CmuA family methyltransferase which potentially enable Sifarchaeota to utilize amide containing compounds and methylated compounds as nitrogen and carbon sources, respectively. Also, a gene encodes for arsenate reductase was potentially acquired from candidate division KSB1 bacterium, potentially allowing Sifarchaeota to use arsenate containing compounds as final electron acceptors.

## Discussion

In this study, we recovered three MAGs from deep Costa Rica sediments belonging to a new Asgard phylum, forming a sister clade to MAGs belonging to Thorarchaeota. We propose the name Sifarchaeota for this phylum, named after the Norse goddess, Sif, wife of Thor. Putative collective metabolic profiles of the Sifarchaeota MAGs showed remarkable differences in the life style and niche adaptations compared to the other Asgard members. We predict a saccharolytic, fermentation-based life style with limited amino acid and fatty acid metabolism, whereas most of the Asgard archaea identified before this study were known for their peptide degradation and short chain fatty acid oxidation capacities (3)(5)(42). We detected genes encoding for incomplete methanogenesis pathways coupled with the carbonyl branch of WL pathway suggesting the capability of Sifarchaeota to perform anaerobic methylotrophy enabling the utilization of various methylated compounds (e.g. methanol and methylamines). The widespread of anaerobic methylotrophy in multiple benthic archaea highlights the importance of this pathway as an effective strategy to utilize various methylated compounds commonly encountered in the marine sediment niches (8)(43). On the other hand, Sifarchaeota MAGs shared similar potential biogeochemical functions with other Asgard archaea including the presence of nitrite reductase (*nirBD*) genes, putatively enabling Sifarchaeota members to reduce nitrite to ammonia as well as genes encoding for sulfate adenylyltransferase (*sat*) and phosphoadenosine phosphosulfate reductase (*cysH*), signifying their putative capability to perform assimilatory sulfate reduction and sulfate activation. These shared characteristics between Asgard genomes confirm the significant roles of different Asgard lineages in nitrogen and sulfur biogeochemical cycles in marine sediment environments.

We also gauged the role of HGT in shaping the evolution of Sifarchaeaota. Our analysis suggests that HGT events either added novel genes to the Sifarchaeota pangenome that impart new functions e.g. butanoate metabolism and biosynthesis of secondary metabolites or enabled the utilization of alternative non-organic compounds as electron source e.g. arsenate reductase. Moreover, we explored the range of horizontally transferred gene donors and we concluded that HGT is not limited to a specific phylogenetic group and probably acquired from the surrounding bacterial communities, normally present in deep marine sediments. Finally, 91% of HGT events are Sifarchaeota lineage specific and probably took place relatively recently during the course of evolution, after the diversification of Sifarchaeota from the other Asgard lineages. We identified only 9% of the events happened earlier to the full diversification of Sifarchaeota from other Asgards, as well as other archaeal lineages. This could be a plausible explanation for the presence of shared functions of non-archaeal origin between most of Asgard lineages.

## Acknowledgements

We would like to thank the shipboard scientists and crew of IODP Expedition 334 for collecting these sediments, and the shorebased curators of the Gulf Coast Repository for their faithful stewardship of precious frozen subsurface samples. We also want to thank our high-performance computation cluster administrator, Karol Miaskiewicz, for his tireless work. This work was supported by the WM Keck Foundation award to JFB.

---

**Supplementary Figure 1.**
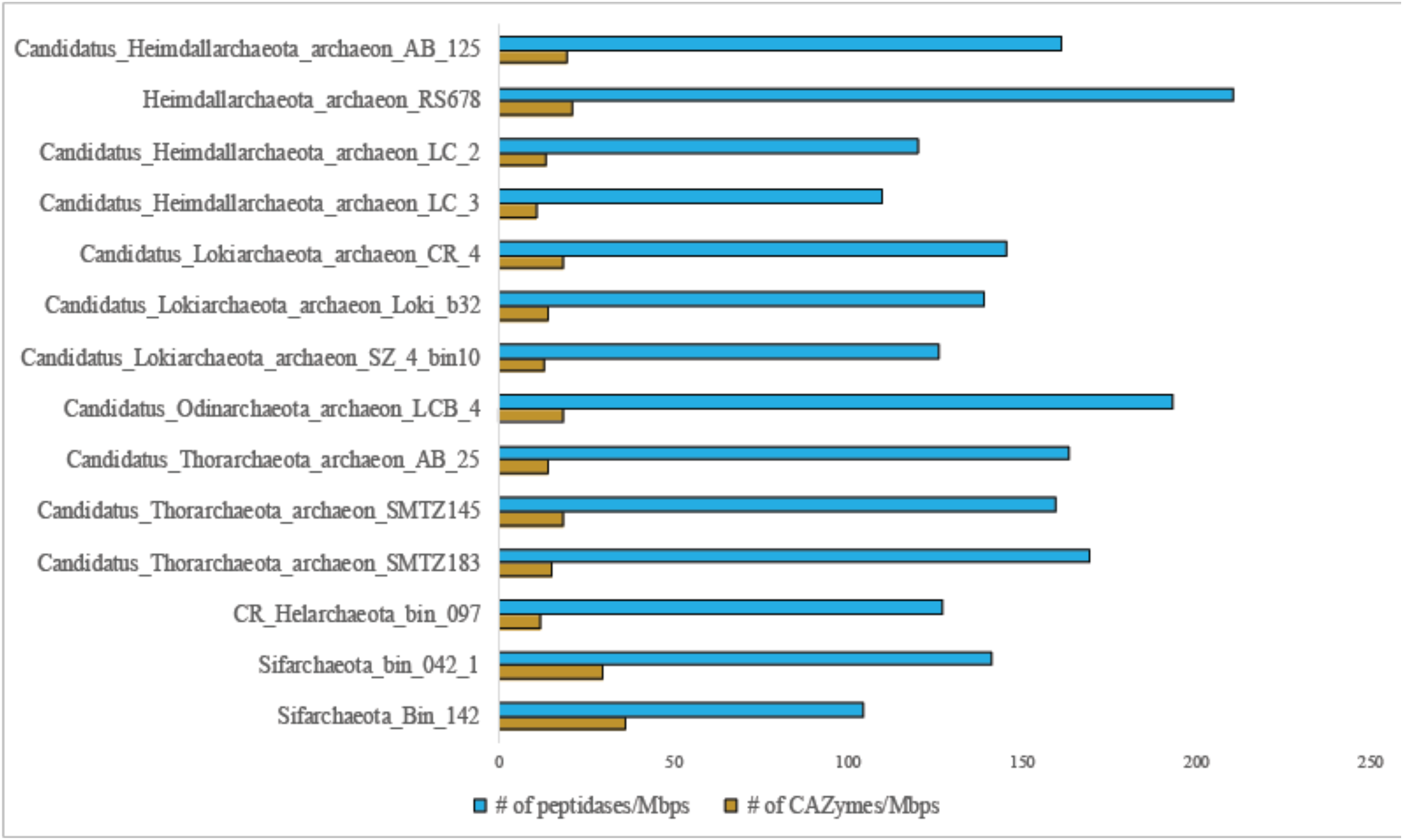
Relative densities (numbers per 1 Mb) of peptidases and CAZymes encoded by the different Asgard MAGs.

**Supplementary Figure 2.**
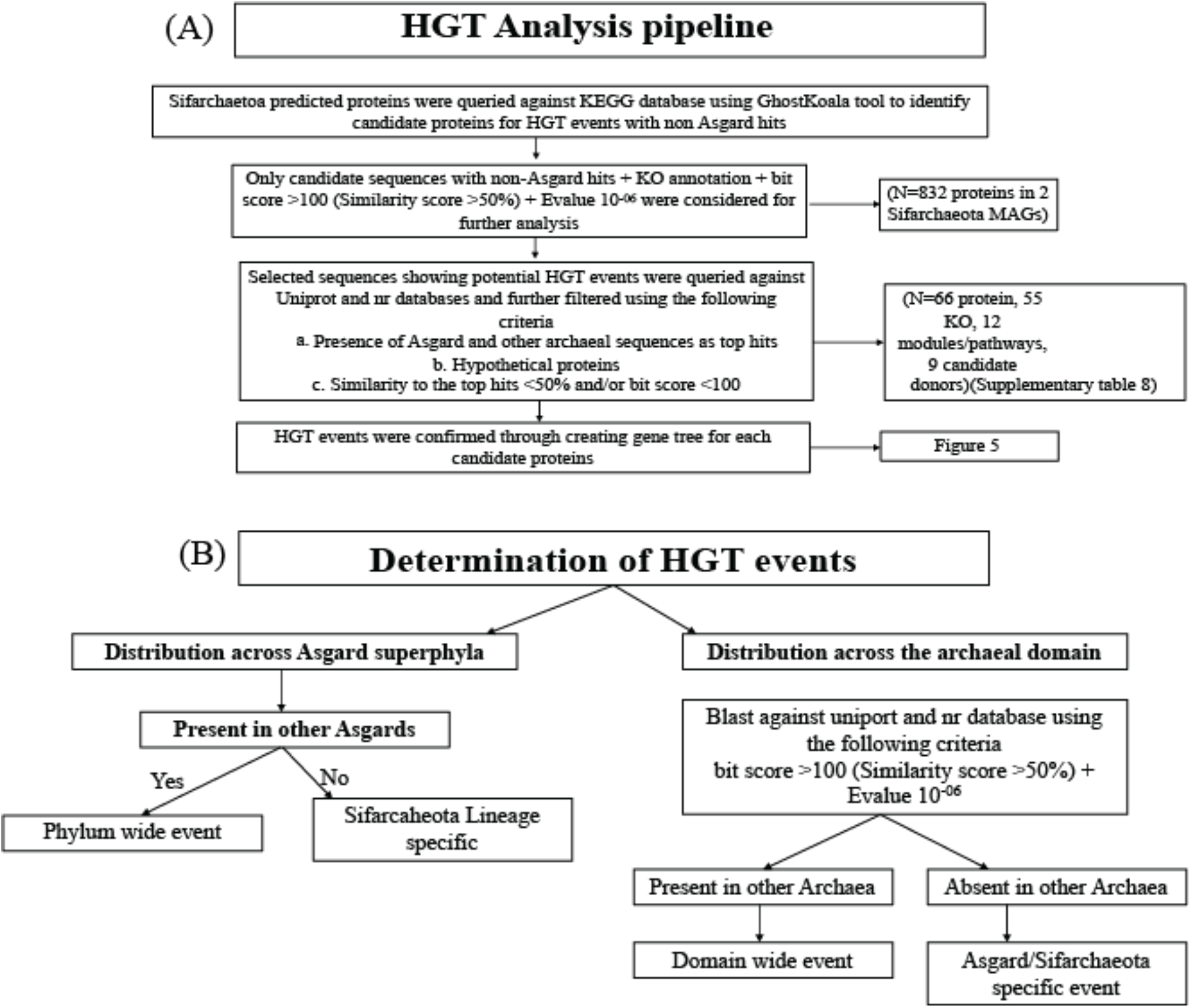
Workflow diagram describes the steps followed to identify HGT events in Sifarchaeota MAGs.

